# Continuous monitoring of deficit irrigation in avocado across two contrasting rainfall years using sensor networks, telemetry, and machine learning

**DOI:** 10.64898/2026.09.03.745400

**Authors:** Alberto Férez-Gómez, Lucía Soler-Escámez, Cristina Ferrer-Blanco, Adrián Pérez-Aguilar, José I. Hormaza, Almudena Díaz-Zayas, Juan M. Losada

## Abstract

Water scarcity and increasingly irregular rainfall threaten avocado production in Mediterranean regions, yet the long term physiological responses of mature trees to sustained deficit irrigation remain poorly understood. We conducted a two year field study integrating continuous monitoring of the soil–plant–atmosphere continuum, drone based multispectral imaging, canopy structural analysis, and AI based fruit phenotyping in a mature avocado orchard subjected to three irrigation regimes. The two years of study differed markedly in rainfall, providing a unique opportunity to evaluate how environmental conditions modulate tree responses to water limitation. Trees under severe deficit irrigation showed depletion of water in deeper soil layers and a flattened physiological profile, with near zero diel variation in leaf thickness and trunk water potential, indicating minimal transpiration and decoupling of tree water status from environmental demand. Drone telemetry via NDVI detected stress during fruit growth and maturation, but not during flowering or the new summer leaf flush, revealing greater drought sensitivity at later maturation stages. Although canopy area did not differ among irrigation treatments, canopy surface roughness increased significantly under deficit irrigation, thereby identifying a novel structural indicator of drought stress. Despite large physiological differences among treatments, fruit number remained stable, while fruit weight decreased significantly under severe deficit irrigation—particularly in the wetter year—suggesting that annual rainfall modulates the trade-off between fruit retention and fruit growth. This study provides the first continuous, multi scale characterization of avocado performance under sustained deficit irrigation in Mediterranean conditions. By integrating plant based sensors, remote sensing, and artificial intelligence, we reveal previously undescribed stress dynamics and identify new indicators for precision irrigation management in fruit crops.

## 1. Introduction

Mediterranean avocado production is increasingly threatened by water scarcity and more irregular rainfall, exposing orchards to recurrent water deficits and highlighting an urgent need for data-driven precision irrigation systems (Junquera et al., 2025; Massaad et al., 2026). However, traditional irrigation research in perennial fruit crops still relies on discrete, manual measurements that provide only snapshots of plant water status and fail to capture diel hydraulic dynamics, carry-over effects among seasons, or the spatiotemporal heterogeneity of mature orchards (Chartzoulakis et al., 2002; Moreno-Ortega et al., 2019, 2021; Durán-Zuazo et al., 2021; Nemera et al., 2021; Cárceles-Rodríguez et al., 2023). While recent advances in low-cost sensor networks, telemetry, high-throughput unmanned aerial vehicle (UAV) imaging, and machine learning now enable continuous monitoring of the soil-plant-atmosphere continuum (SPAC) (Carella et al., 2024; Velázquez-Chavez et al., 2024), these precision tools are rarely integrated into a unified monitoring framework. Therefore, integrated orchard-scale systems combining continuous plant sensing, multi-depth soil monitoring, canopy imaging, and artificial intelligence to support operational irrigation decision-making remain largely unexplored (Sharma and Shivandu, 2024).

Environmental and soil moisture sensors are widely adopted to guide irrigation decisions (Yin et al., 2021). However, they estimate plant water status indirectly and can easily fail to capture the spatial heterogeneity of the root zone (Wheeler et al., 2023). To address soil limitations, plant-based sensors are also frequently utilized, predominantly focusing on trunk diameter and sap flow (Ortuño et al., 2010). Yet these tools have inherent limitations: sap flow measurements often track evaporative demand more closely than actual water deficit (Wheeler et al., 2023), and trunk diameter fluctuations can lag rapid real-time changes in trunk water potential (Ψ_trunk_) (Blanco and Kalcsits, 2023). Although advanced continuous sensors, such as ZIM probes for leaf turgor monitoring and microtensiometers for Ψ_trunk,_ have expanded the range of physiological measurements available, their deployment remains restricted to a small number of tree crops (Rodríguez-Domínguez et al., 2019). Recent studies emphasize that single-sensor approaches miss complex, cultivar-specific drought strategies (Marino et al., 2021). Furthermore, accurately capturing diel water uptake dynamics requires integrating multi-depth soil moisture profiles (Calabritto et al., 2024; Magh et al., 2026) with atmospheric demand (Pagay, 2021). Consequently, while the physiological responses of individual trees are well documented, the combined use of continuous Ψ_trunk_, leaf thickness dynamics, and multiple-depth soil moisture has rarely been evaluated in mature orchards. As a result, how plant hydraulics coordinate with soil water availability and atmospheric demand under sustained deficit irrigation remains poorly resolved, constraining the development of robust, orchard-scale precision irrigation strategies.

Remote sensing has been increasingly used to assess canopy vigor and water status in perennial fruit tree crops (Campos et al., 2021). However, applications in avocado have largely focused on Normalized Difference Vegetation Index (NDVI) using coarse or infrequent time series (Robson et al., 2017). While NDVI varies with water stress across phenological stages, its sensitivity depends strongly on the growth stage, canopy architecture, and sensor resolution. For instance, studies in other tree crops, such as almonds, have shown that NDVI differences became apparent only under moderate water stress and were cultivar-dependent, limiting the use of NDVI for early-stage stress detection (Gutiérrez-Gordillo et al., 2021). Furthermore, while fractional cover and canopy size indicate an apparent response to water deficit, structural metrics alone are rarely validated against chronic tree water stress (Berry et al., 2025). Moreover, traditional two-dimensional (2D) metrics such as projected canopy area fail to capture vertical thinning, internal defoliation, or microstructural degradation. These limitations highlight the need for robust, high-resolution structural assessments, such as AI-based canopy segmentation and canopy surface roughness (CSR), to accurately evaluate long-term irrigation responses.

The impact of water scarcity on avocado yield remains contradictory. Sustained deficit irrigation can significantly reduce both total yield and fruit size (Durán-Zuazo et al., 2021), but moderate water reductions have shown minimal effects on crop load and overall production under specific conditions (Beyá-Marshall et al., 2022). Furthermore, while technological approaches have improved, comprehensive integrations remain insufficient. For example, although recent studies have combined remote sensing and soil moisture sensors to optimize avocado irrigation management, most lack multi-year yield-validation frameworks that link physiological responses to crop productivity (Torres-Quezada et al., 2025; Kaneko et al., 2026). Similarly, while remote sensing has been shown to improve yield prediction in several perennial fruit-tree models (Bai et al., 2019), these methods have not been validated through empirical field studies in avocado. As emphasized in recent reviews, reliable horticultural yield sensing requires combining multiple complementary technologies (Longchamps et al., 2022). Even so, to date, no study has fully integrated multi-year yield assessments with field-based physiological and remote sensing monitoring under different drought conditions.

Finally, while the use of artificial intelligence for fruit phenotyping is advancing rapidly, its application remains irregular across traits and crops. Although deep learning has become widely adopted for fruit detection and counting, estimation of precise physical traits, such as fruit weight, is limited to a few species (Miranda et al., 2023). In avocado, current deep learning applications have largely focused on basic canopy detection and postharvest ripeness assessment, leaving a relevant gap for robust models capable of precise fruit weight estimation from 2D imagery (Davur et al., 2023). Moreover, abiotic stress phenotyping is generally underrepresented in AI-driven agricultural studies (Houetohossou et al., 2023). Therefore, developing an automated computer vision approach to accurately estimate fruit weight could significantly optimize high-throughput evaluation of crop productivity and yield components across different water-deficit regimes.

This study presents a two-year field experiment in a mature ‘Hass’ avocado orchard subjected to three irrigation regimes under contrasting annual rainfall conditions. We deployed a 5G-enabled sensor network to collect continuous Ψ_trunk_, leaf thickness, and soil volumetric water content (VWC) at multiple depths. In parallel, we conducted weekly UAV multispectral and photogrammetric surveys to monitor NDVI and extract aerial canopy metrics using YOLO-based instance segmentation. Additionally, we developed a separate YOLO pipeline for precise fruit phenotyping, processing avocados photographed individually under controlled laboratory conditions. Our objectives were to (i) characterize diel and seasonal hydraulic responses to sustained deficit irrigation, (ii) evaluate the sensitivity of spectral (NDVI) and structural (canopy area and CSR) UAV metrics across different irrigation treatments and/or phenological stages, and (iii) develop and validate an AI model for automated fruit weight estimation. We hypothesized that severe water deficit would reduce diel physiological performance, indirectly showing stomatal closure and hydraulic decoupling, and that NDVI responses to irrigation would be highly dependent on the tree’s phenological stage. By integrating continuous plant sensors, telemetry, UAV remote sensing, and machine learning, this study provides a comprehensive framework for characterizing avocado responses to sustained deficit irrigation and advances the development of a practical, data-driven framework for precision irrigation in perennial orchards.

## 2. Materials and Methods

### 2.1. Experimental site and 5G network deployment

The field trial was conducted at the Institute for Mediterranean and Subtropical Horticulture “La Mayora” (IHSM) in Málaga, Spain, within a 50-ha experimental station dedicated to the study of subtropical crops. The study was performed in a mature avocado orchard (*Persea americana* Mill., cv. Hass) with an area of 9,502 m2, coordinates 36° 45 26.67 “ N 4° 02’ 26.46” W, and an elevation of 68 m above sea level (Fig. 1A).

**Figure 1.**
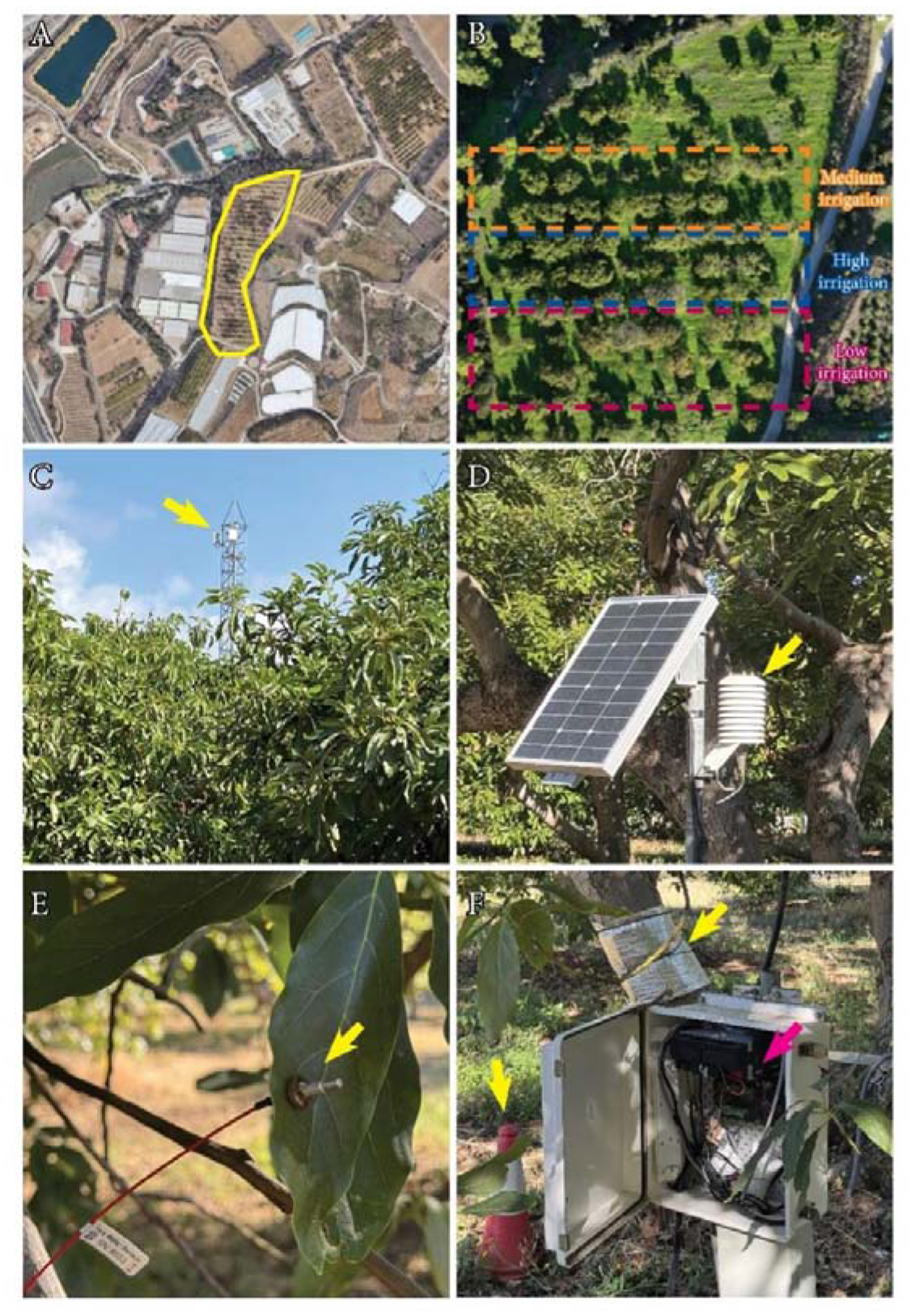
Experimental design. (A) Location map of the avocado field within the Experimental Station of the Institute of Subtropical and Mediterranean Hortofruticulture (IHSM-CSIC-UMA) 36°45’26.67“ N 4°02’26.46” W. (B) Drone RGB orthomosaic showing the three irrigation tree lines (dotted). (C) Telecommunication tower with a 5G antenna (yellow arrow) used to transmit data from the field data logger to the local server. (D) Climate station (yellow arrow) for temperature and humidity. (E) Leaf patch clamp sensor (yellow arrow). (F) Trunk water potential sensors (yellow arrow, top) are protected with a gray insulating mat; the cabinet with the battery for the data logger (pink arrow) collects sensor signals, including the soil volumetric water content sensor protected by the cone (yellow arrow, left).

To ensure that the experiment reflected commercial field conditions, mature’Hass’ avocado trees (>40 years old) were selected, intentionally incorporating the natural structural and architectural heterogeneity typical of mature orchards. In early 2023, the orchard was structured into three distinct irrigation treatments. Each treatment was applied to at least two adjacent tree rows, with at least seven trees per row. The standard historical schedule for the orchard was maintained as the fully irrigated control, designated as the ‘High’ irrigation treatment (100% of crop water requirement). The two sustained deficit irrigation treatments received 66% (‘Medium**’**) and 33% (‘Low’) of the control irrigation volume (Fig. 1B). Irrigation events occurred simultaneously across all treatments, with the differential water application achieved by varying the number of active drip lateral lines per tree row.

Finally, to support the high-frequency, real-time data transmission required for continuous environmental and physiological monitoring, a dedicated private 5G network was deployed. Two 5G sites provide coverage to the experimental farm. A macro cell (see Fig. 1C) was installed in an adjacent plot to ensure full coverage over the avocado experimental site without interfering with the orchard’s canopy architecture. This cell radiates in band 40, which operates in the 2300–2400 MHz range under a Time Division Duplex (TDD) configuration. It is a reasonable candidate for providing wireless network coverage to agricultural sites if the farm is compact-to-medium-sized and high data throughput is a priority. Band 40 also provides good performance for delivering video streaming from cameras and drones, backhauling IoT sensor data, providing a real-time monitoring dashboard, and remotely controlling the drones. The 5G private network is connected to the Internet via satellite, providing global connectivity and the ability to offer third-party real-time access to the data and images collected.

### 2.2. Sensor network and continuous physiological monitoring

To monitor the orchard microclimate, environmental sensors were installed on a representative tree within the ‘Low’ irrigation regime to continuously track ambient temperature and relative humidity (Fig. 1D). These data were used to calculate atmospheric vapor pressure deficit (VPD). First, the saturation vapor pressure (*e*_s_, in kPa) was estimated using the Tetens equation:

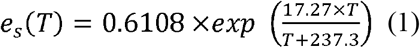

Where *T* is the temperature in degrees Celsius (°C). Subsequently, VPD was calculated as:

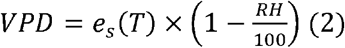

Where *R H* represents the relative humidity (%).

To monitor the soil-plant continuum, a specific subsample of trees was sensorized within each irrigation treatment. Soil capacitance sensors (Sentek PLUS, Sentek Sensor Technologies, Stepney, SA, Australia) were installed to continuously track VWC at three depths: 10 cm, 30 cm, and 50 cm. Inside each treatment row, these soil probes were distributed across three consecutive trees and placed at a consistent distance from the sprinklers to minimize spatial variability.

Among these three trees, one representative tree per irrigation treatment was additionally equipped with plant-based sensors to monitor diel hydraulics. Two microtensiometers (FloraPulse, Davis, CA, USA) were inserted into the trunk of each representative tree. Following the manufacturer’s protocols, the sensors were covered with insulating, reflective mats to protect against thermal fluctuations and wildlife. Additionally, at least one magnetic leaf patch pressure probe (ZIM Plant Technology GmbH, Hennigsdorf, Germany) was installed on a fully expanded, healthy leaf representative of the same tree (Fig. 1E) to continuously monitor changes in leaf turgor. Following calibration, data were obtained in mV and were transformed into units of pressure (p_p_, in kPa) using the manufacturer instructions.

To integrate all parameters in real time, the sensors were connected to a data logger (Campbell Scientific) programmed to record measurements at 15-minute intervals throughout the experimental period (Fig. 1F). The data loggers were further linked to a 5G router, enabling high-frequency telemetry to a local server. All time-series data were systematically stored in InfluxDB and processed for real-time visualization on an open-access Grafana dashboard, enabling continuous public monitoring of the orchard’s hydraulic status (http://150.214.47.156:43000/dashboards/f/dds7hz0x3ow74b/).

### 2.3. UAV remote sensing and canopy segmentation

To evaluate the structural canopy area of the avocado trees, weekly flights were conducted in 2025 using a DJI Matrice 600 drone equipped with an RGB camera (DJI Zenmuse Z3) and 5G connectivity to deliver images to the edge server hosting the AI algorithm. Flights were conducted at an altitude of 30m above ground level, with continuous video recorded from which individual images were extracted at a rate of 1 frame per second. A dataset of 775 frames was manually annotated using Roboflow software (Dwyer et al., 2026) to train a YOLOv11n-seg instance segmentation model. During the inference phase, a confidence threshold of 0.5 was established to ensure robust mask predictions. Finally, a custom Python script was developed to translate the pixel-based segmentation into physical metrics, calculating the estimated true canopy area using the Ground Sample Distance (GSD) conversion factor for the 30 m flight altitude.

For spectral evaluations, specifically the NDVI, a DJI Mavic 3 Multispectral (Mavic 3M) drone was utilized. The UAV is equipped with an integrated incident light sensor and multispectral imaging system including four 5 MP bands: Green (560±16 nm), Red (650±16 nm), Red Edge (730±16 nm), and Near-Infrared (NIR) (860±26 nm). Weekly flights were conducted from December 2025 through May 2026 at an approximate altitude of 60 m, yielding a GSD of 3 cm/pixel.

### 2.4. Photogrammetry and real-time spatial data analysis

Following data acquisition, individual RGB and multispectral images were processed using OpenDroneMap (ODM) software to generate dense point clouds and orthomosaics (RGB and multispectral). Radiometric calibration was applied to the multispectral dataset using metadata from the drone’s incident light sensor to compensate for variations in solar irradiance across flights. To ensure high-fidelity data extraction and to strictly isolate the trees within each irrigation regime from background elements, the RGB orthomosaic was imported into QGIS, where tree canopies for the three irrigation treatments were manually delineated using precise vector polygons. Concurrently, the multispectral orthomosaic was processed in R (R Core Team (2024)) using the *terra* package (Hijmans R (2025), version 1.8-50) to calculate the NDVI at the pixel level according to the standard formula:

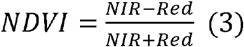

To isolate the true canopy reflectance and eliminate background noise, such as soil or shade, an empirical threshold was applied to exclude pixels with NDVI values below 0.5. Finally, the vector polygons defined in QGIS were used as zonal masks to extract the mean NDVI value for each tree canopy for subsequent statistical analysis.

For the three-dimensional structural analysis of avocado trees, CSR, the generated dense point clouds were processed using the *lidR* (Jean-Romain Roussel and David Auty, 2026), version 4.2.3, and *terra* (Hijmans R, 2025), version 1.8-50, packages in the R statistical environment. First, a Digital Surface Model (DSM) was created at a spatial resolution of 0.1 m by applying a point-to-raster algorithm. Then, a Digital Terrain Model (DTM) was generated at the same spatial resolution using exclusively the points previously classified as ground by the ODM photogrammetric engine. This was achieved by applying a k-nearest neighbors (k = 10, p = 2) Inverse Distance Weighting (IDW) interpolation algorithm.

The Canopy Height Model (CHM), which represents the true or orthometric height of the vegetation above ground level, was obtained through a pixel-by-pixel subtraction of the DTM from the DSM (CHM = DSM – DTM). Any resulting negative values, typically associated with photogrammetric noise, were reassigned to zero. Finally, the CHM was spatially aligned with the previously mentioned vector polygons delineating individual tree crowns. To evaluate the effects of different irrigation levels on the crown’s internal architecture, zonal statistics were extracted at the individual-tree level using the *terra* package. The CSR parameter was calculated as an indicator of structural homogeneity and foliage density. This CSR is defined as the standard deviation of the heights of all CHM pixels falling within the bounding polygon of each crown. Low values indicate a dense, continuous, and compact canopy (reduced variation in outer foliage surface heights), while high values represent an irregular canopy with partial defoliation.

To automate these processes, an algorithm for real-time transmission was developed, comprising a drone-based system equipped with a Pixhawk V6X flight controller, a Raspberry Pi, and an onboard 5G modem. Telemetry and control data were forwarded through the Raspberry Pi to a ground station running the Mission Planner application, enabling bidirectional communication over the private 5G network. Simultaneously, an Insta360 camera streamed RTSP video to a MediaMTX server and, using a modified SDK, captured images every 5 s. These images were automatically transferred to the processing server, where the YOLOv11n-seg model performed canopy segmentation, object detection and counting, and canopy area estimation throughout the programmed flight mission.

### 2.5. Fruit harvest and computer vision-based weight estimation

Yield parameters were evaluated across two consecutive productive cycles. During the 2024-2025 campaign (harvested in spring 2025), total fruit count and fruit weight were recorded per tree. In the 2025-2026 campaign, to ensure an accurate evaluation of total productivity, the harvest included all fruits remaining on the trees, as well as those that had fallen before harvest due to severe winter winds (February 2026).

In addition to recording the total fruit count, a representative subset of 1004 individual avocado fruits, from the 2025-2026 campaign, was selected to estimate their weights using computer vision. Each fruit was individually weighed on a precision balance and photographed under controlled laboratory lighting using a Nikon D3500 (Nikon AF-P DX NIKKOR 18-55mm f/3.5-5.6g VR camera lens) mounted at a fixed distance of 50 cm. This image dataset was used to train a YOLOv8s-seg instance segmentation model to accurately extract the 2D projected area (*A*, measured in cm^2^) of each fruit. An empirical quadratic regression model was then derived to predict actual fruit mass based on the segmented area:

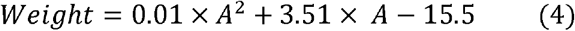

### 2.6. Data processing and statistical analysis

Time-series data recorded at 15-minute intervals were retrieved from the central database (InfluxDB) and preprocessed in Python using pandas (McKinney, 2010, version 1.2.4) and NumPy (Harris et al., 2020, version 1.20.1), following a common quality-control workflow for the three irrigation treatments. Complementary records associated with the same timestamp were merged, and the analysis was restricted to the period from 22 April 2024 to 21 April 2026. Duplicate timestamps were subsequently resolved, and the series from the three treatments were aligned to a regular 15-minute time grid.

Redundant channels were evaluated based on their temporal coverage, the plausibility of their readings, and the temporal coherence of the signal. Observations outside the reference ranges established for each sensor type—air temperature, −10 to 55 °C; relative humidity and soil volumetric water content, 0 to 100%; leaf patch signal, 3 to 100 mV/V; and trunk water potential, −3 to 0 MPa—were flagged as missing values. At each timestamp and depth, the available readings from the three soil probes installed in each treatment were averaged to obtain a single series per treatment and depth.

For the seasonal analysis (Fig. 2), the cleaned 15-minute series were summarized at a daily resolution using maximum VPD, minimum trunk water potential, and mean soil volumetric water content at each depth. Daily rainfall records were obtained from the official National Agency of Meteorological Station (AEMET 6201X), located at the Experimental Station, and incorporated at their original daily resolution. On days with less than 50% coverage of the local meteorological variables, daily climate statistics were gap-filled using AEMET records when all required variables were available. This procedure was applied exclusively to the daily aggregates and not to the 15-minute series. Daily leaf patch pressure amplitude was calculated as the difference between the daily maximum and minimum only when at least 50% of the expected records were available—48 of the 96 daily observations— and was converted to kPa using the sensor transformation constant (K = 80). Days showing abrupt shifts in the overall signal level or amplitudes inconsistent with their local temporal dynamics, suggestive of possible repositioning of the leaf probe, were flagged as anomalous. Internal gaps comprising one or two consecutive missing observations, and bounded by valid observations, were linearly interpolated. Longer interruptions were retained as missing values to avoid introducing artificial patterns.

**Figure 2.**
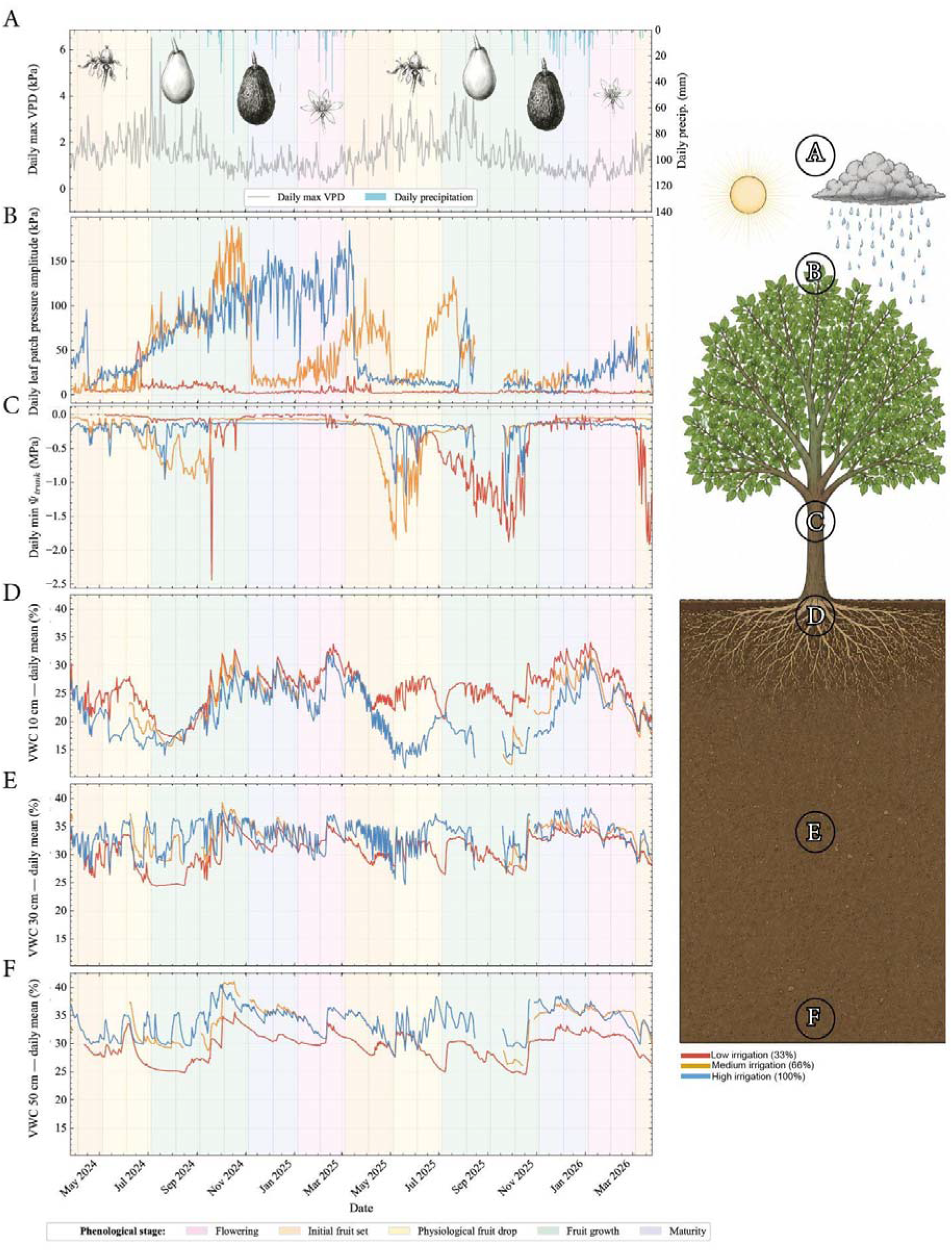
Seasonal dynamics of the soil-plant-atmosphere continuum across phenological stages from April 2024 to April 2026. Colored backgrounds indicate the phenological stages labeled within the figure, also pictured with drawings of fruits. (A) Daily maximum vapor pressure deficit (VPD; grey line) and daily precipitation recorded by the AEMET 6210X meteorological station (blue bars). (B) Daily leaf patch pressure amplitude (kPa). (C) Daily trunk water potential (MPa). (D-F) Daily mean soil volumetric water content at a 10cm depth(D), 30cm depth (E), and 50 cm depth (F). Treatments represented by colors: blue (high or 100% field capacity), orange (medium or 66%), and red (low or 33%). Color-coded phenological stages (bottom legend): light pink (flowering), yellow (initial fruit set), light green (fruit growth), light blue (maturation).

For the diel analysis (Fig. 3), the cleaned series were retained at their original 15-minute resolution during August 2024, selected as the month of highest atmospheric demand of 2024, during fruit growth. The leaf patch pressure signal was plotted as a continuous 15-minute time series rather than summarized as a daily amplitude and was expressed in kPa using the same constant (K = 80). Daily rainfall from the AEMET station was overlaid at its original daily resolution, and day and night periods were delineated using local sunrise and sunset times. Both figures were generated in Python using Matplotlib (Hunter, 2007, version 3.3.4). Subsequent data wrangling and temporal filtering for the statistical analyses were performed in R using the *dplyr* (Wickham et al., 2023, version 1.1.4) and *lubridate* (Grolemund and Wickham, 2011, version 1.9.4) packages. Remaining missing values or technical anomalies resulting from sensor maintenance or connection loss were treated as NA and excluded from subsequent analyses.

**Figure 3.**
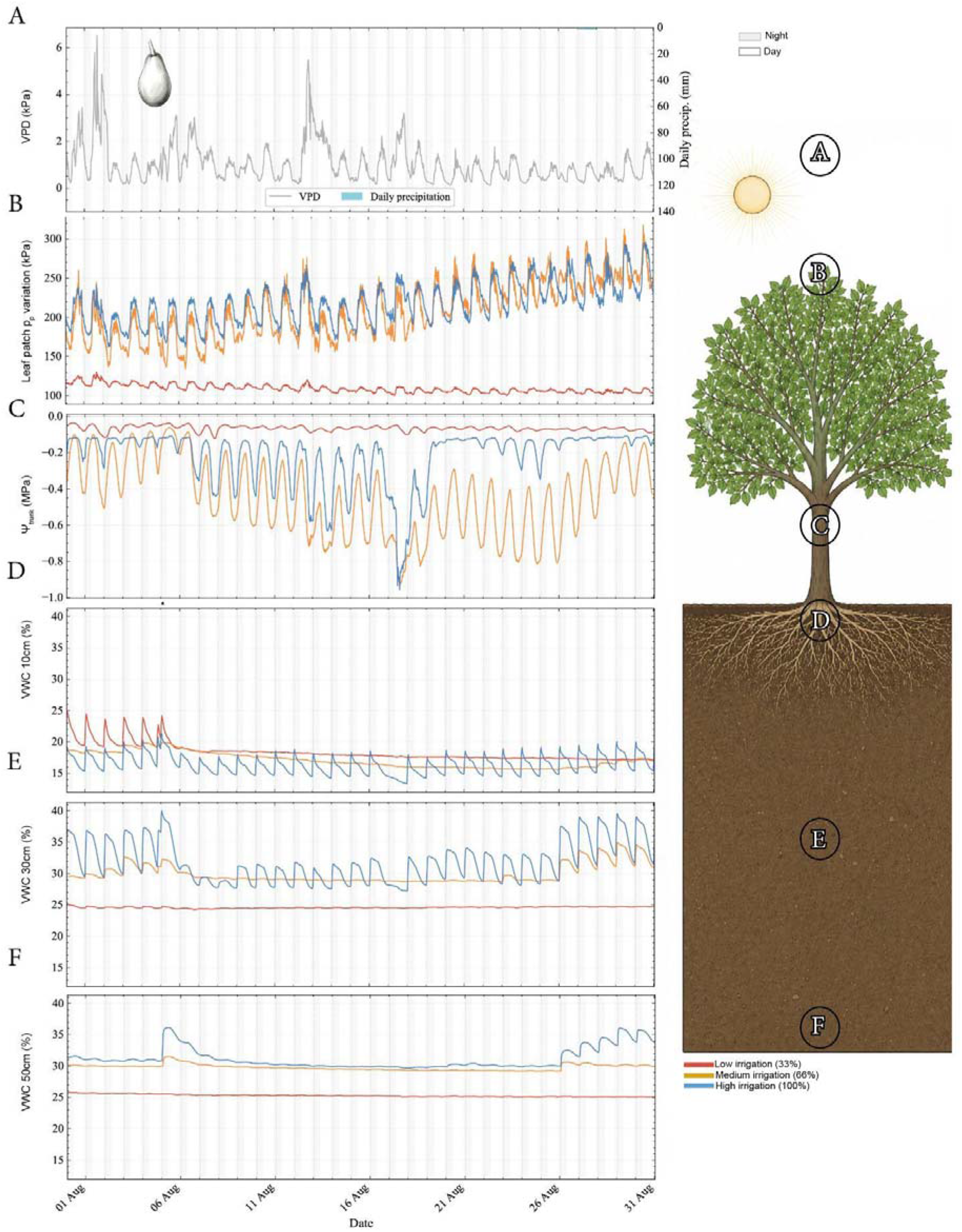
Diel dynamics of the soil-plant-atmosphere continuum during August 2024, the month of highest atmospheric demand within the fruit growth period. White and grey backgrounds indicate day and night periods, respectively. (A) Vapor pressure deficit (VPD; grey line) at a 15-min resolution (kPa) and daily precipitation recorded by AEMET 6201X meteorological station (blue bars) (B) Leaf patch pressure (kPa), showing daytime maxima and nocturnal minima. (C) Trunk water potential showing daytime minima and nocturnal maxima (MPa). (D-F) Soil volumetric water content at a 10cm (D), 30cm (E), and 50 cm depth (F). Irrigation treatments represented by colors: blue (high or 100% field capacity), orange (medium or 66%), red (low or 33%).

All statistical evaluations were conducted using the *stats* (version 4.4.1), *car* (Fox and Weisberg (2019), version 3.1.5), and *ggsignif* (Ahlmann-Eltze and Patil (2021), version 0.6.4) packages in R. Before the global test, the datasets within each irrigation treatment (High, Medium, and Low) were evaluated for parametric assumptions. Normality was assessed using the Shapiro-Wilk test, and homoscedasticity was assessed using Bartlett’s test for normally distributed data or Levene’s test when non-normality was detected.

For datasets that fitted both parametric assumptions, a one-way Analysis of Variance (ANOVA) followed by a pairwise Student’s t-test was carried out. In cases with a normal distribution but without homoscedasticity, Welch’s ANOVA, combined with a pairwise Welch’s t-test, was used. For datasets without a normal distribution, the nonparametric Kruskal-Wallis test was used, followed by the pairwise Wilcoxon post hoc test. P-values were adjusted using the Holm-Bonferroni correction.

## 3. Results

### 3.1. Seasonality and water dynamics in the soil-plant-atmosphere continuum

The phenological cycle of avocado is primarily defined by its reproductive development, which spans approximately one year. During March and April, the trees lose most of their old leaves concomitant with massive flower development and bloom. In June, the initial fruit set is followed by a massive physiological drop in fruitlets. The remaining fruits developed from June to November, with physiological maturity (the availability of ripening after harvest) reached around December. Because avocado is a climacteric fruit, commercial harvest can extend for several months, as fruits remain attached to the tree without ripening and only begin to ripen after detachment. In this study, fruits were harvested from February to March, coincident with the onset of the subsequent flowering cycle (see also Alcaraz et al., 2013).

Both 2024 and 2025 were characterized by VPD maxima in the summer, when rainfall was near zero, and coincided with periods of fruit growth (Fig. 2A). The summer of 2024 followed several consecutive years of drought, and VPD reached a maximum of > 6kPa (Fig. 2A). In contrast, abundant rainfall during autumn and winter 2024-2025 reversed the prolonged drought, making 2025 the wettest year of the decade, interrupting a prolonged hyper-arid cycle. Consequently, maximum VPD in the summer of 2025 was below 4kPa.

Continuous monitoring of leaf thickness revealed contrasting hydraulic responses among irrigation treatments (Fig. 2B). The daily amplitude of the leaf patch p_p_ remained minimal in the tree with low irrigation treatment in both years, except during the initial fruit set period of the fruit set, suggesting a minimum variation (Fig. 2B). In contrast, this parameter was positive and followed a similar pattern in leaves under medium and high irrigation treatments. Throughout 2024, the amplitude of the leaf patch p_p_ increased progressively during fruit growth, reaching maximum values before fruit maturation in the medium irrigation treatment and during fruit maturation in the high irrigation treatment, then declining during leaf senescence. In the wet year 2025, leaf patch p_p_ dynamics exhibited comparable seasonal patterns in all treatments.

Interestingly, these leaf-level dynamics mirrored the variations in the water potential of the stem, or Ψ_trunk_ (Fig. 2C). For example, the Ψ_trunk_ values remained stable and close to 0.0 MPa in the trees with the low-irrigation treatment throughout 2024, suggesting low tension in the xylem. In contrast, trees under medium and high irrigation treatments showed more negative Ψ_trunk_ values, reaching seasonal minima of approximately-1.0 MPa during the fruit growth period. During the wet year 2025, all irrigation treatments showed negative Ψ_trunk_ values, but asynchronously: medium-and high-irrigation treatments reached their lowest potentials during the initial fruit set, whereas low irrigation trees delayed their drop to negative Ψ_trunk_ values until the fruit maturity phase in autumn.

In the soil profile, the VWC in the most superficial layer was similar among all three irrigation regimes (Fig. 2D), oscillating from a minimum of 15% during the fruit growth periods of both years to a maximum of 30-35% during maturation and flowering (coincident with the period of rains). This baseline pattern was conserved in the intermediate soil layer for all three treatments (Fig. 2E). However, in the deepest measured layer, the low irrigation treatment reached a minimum of 25% during the dry summer of 2024 (Fig. 2F) compared to higher VWC values sustained by the medium and high irrigation treatments. This vertical gradient was maintained throughout the two-year study period, with greater depletion in the deeper soil layers under the restricted-irrigation treatment.

### 3.2. Diel water dynamics in the continuum soil-plant-atmosphere

To evaluate the high-resolution temporal perception provided by the agrotechnical sensors, we examined a representative month in detail: August, the hottest and driest month of 2024. This month was characterized by a single minor rainfall event and extreme fluctuations of VPD (Fig. 3A). The leaf patch p_p_ oscillated during the diel cycle, with minimum values at night and maximum values during the day (Fig. 3B). Comparison among irrigation treatments revealed that the amplitudes of diel variations were larger in medium and high irrigation treatments compared with the minimal variations exhibited by the leaves under low irrigation treatment (Fig. 3B). Similarly, the Ψ_trunk_ showed diel variations in the potential registered (Fig. 3C), yet the variation was minimal for the trees under low irrigation treatment (Fig. 3C). In contrast, the Ψ_trunk_ of the trees under medium and high irrigation treatments exhibited more negative potentials, with minima around-1.0 MPa (Fig. 3C). In the soil matrix, sensors of the superficial layers captured individual irrigation events among all treatments, fluctuating between 15 and 25 % (Fig. 3D). However, in the intermediate (Fig. 3E) and deeper soil layers (Fig. 3F), dynamic variations and water refilling were only observed under the medium and high irrigation treatments. In these deeper layers, the low irrigation treatment maintained a consistently lower water percentage.

### 3.3. Remote sensing and computer vision for avocado canopy characterization

Weekly UAV flights were used to train a YOLOv11n-seg algorithm to automatically define the tree canopy area (Fig. 4A), achieving a mean Average Precision (mAP@50) of 89%. Interestingly, no significant differences were found in the tree canopy area among the three different irrigation treatments, demonstrating that 2D canopy projection area alone is a poor indicator of drought stress in mature avocado orchards due to high tree-to-tree morphological heterogeneity (Fig. 4B). On the other hand, the CSR successfully discriminated against irrigation treatments (Fig. 4C). Trees under the low irrigation treatment showed significantly higher CSR values compared with trees under medium and high irrigation treatments, which displayed no significant differences among them. This structural alteration implies that severe water deficit triggers partial defoliation and formation of foliar gaps in the canopy (Fig. 4D).

**Figure 4.**
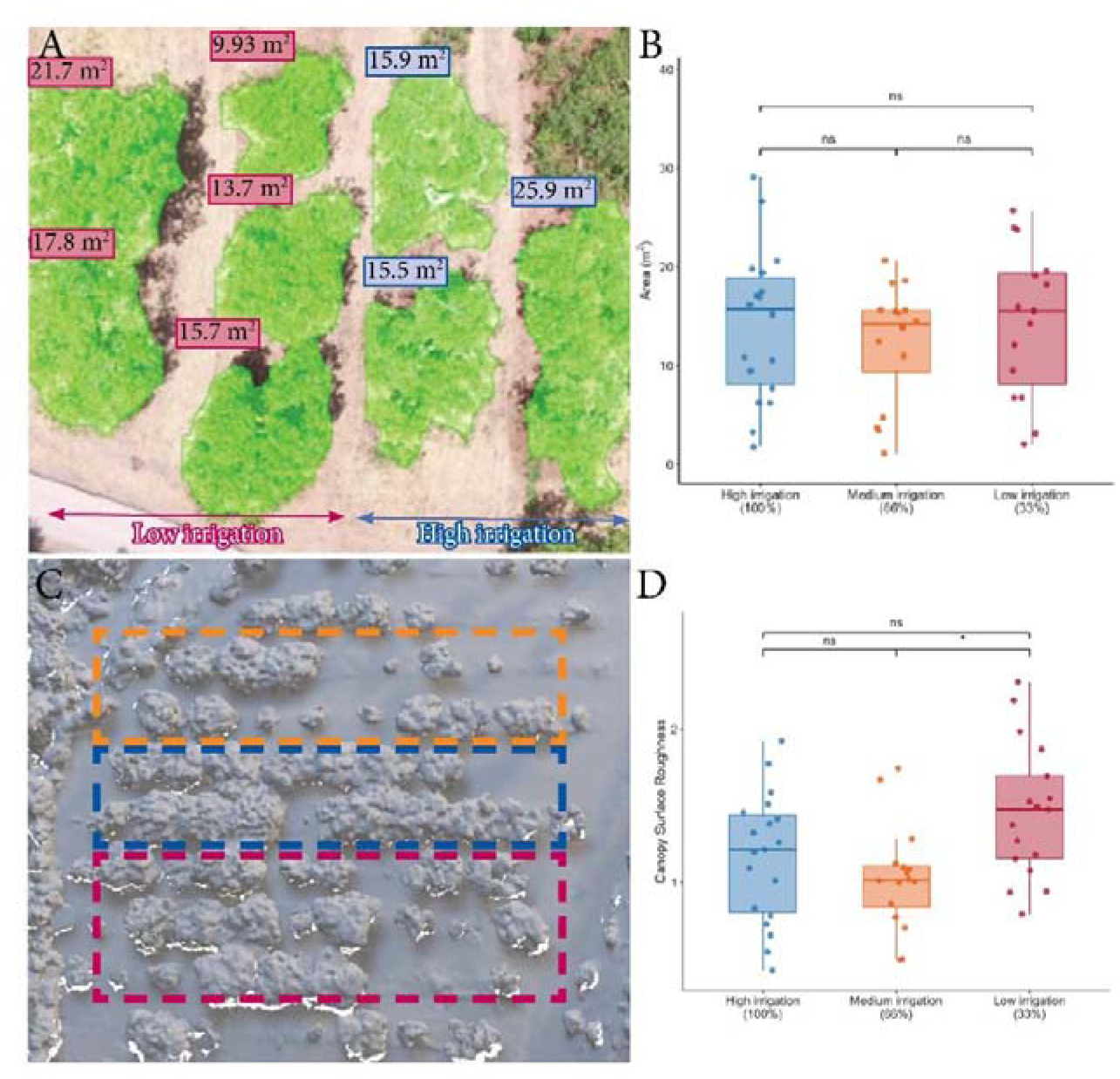
Canopy telemetry and remote sensing-derived canopy metrics. (A) YOLO inference example for avocado canopy segmentation and real-time measurement of canopy area. (B) Boxplot displaying canopy area (m^2^) by irrigation treatment (High: blue; Medium: orange; Low: red). (C) 3D point cloud derived from drone imagery; dotted lines indicate the irrigation treatments. (D) Canopy surface roughness (CSR) by irrigation treatment. ANOVA test followed by pairwise *t*-tests with Holm’s correction; significance comparing irrigation levels at *p* < 0.05(*), or “ns” (not significant).

NDVI maps were generated during the latest fruit maturation phenological stages. Through the fruit maturation stage (Fig. 5A), while no differences were quantified between the medium and high irrigation treatments, significant differences in the NDVI index were depicted for the trees under low irritation treatment (Fig. 5B). These differences were maintained across the months of fruit development and harvest, when the trees kept the leaves from the previous year (Fig. 5C). However, at the time of new flush and flowering (Fig. 5D), which coincided with the seasonal winter rains, the NDVI values showed no significant differences across treatments (Fig. 5E).

**Figure 5.**
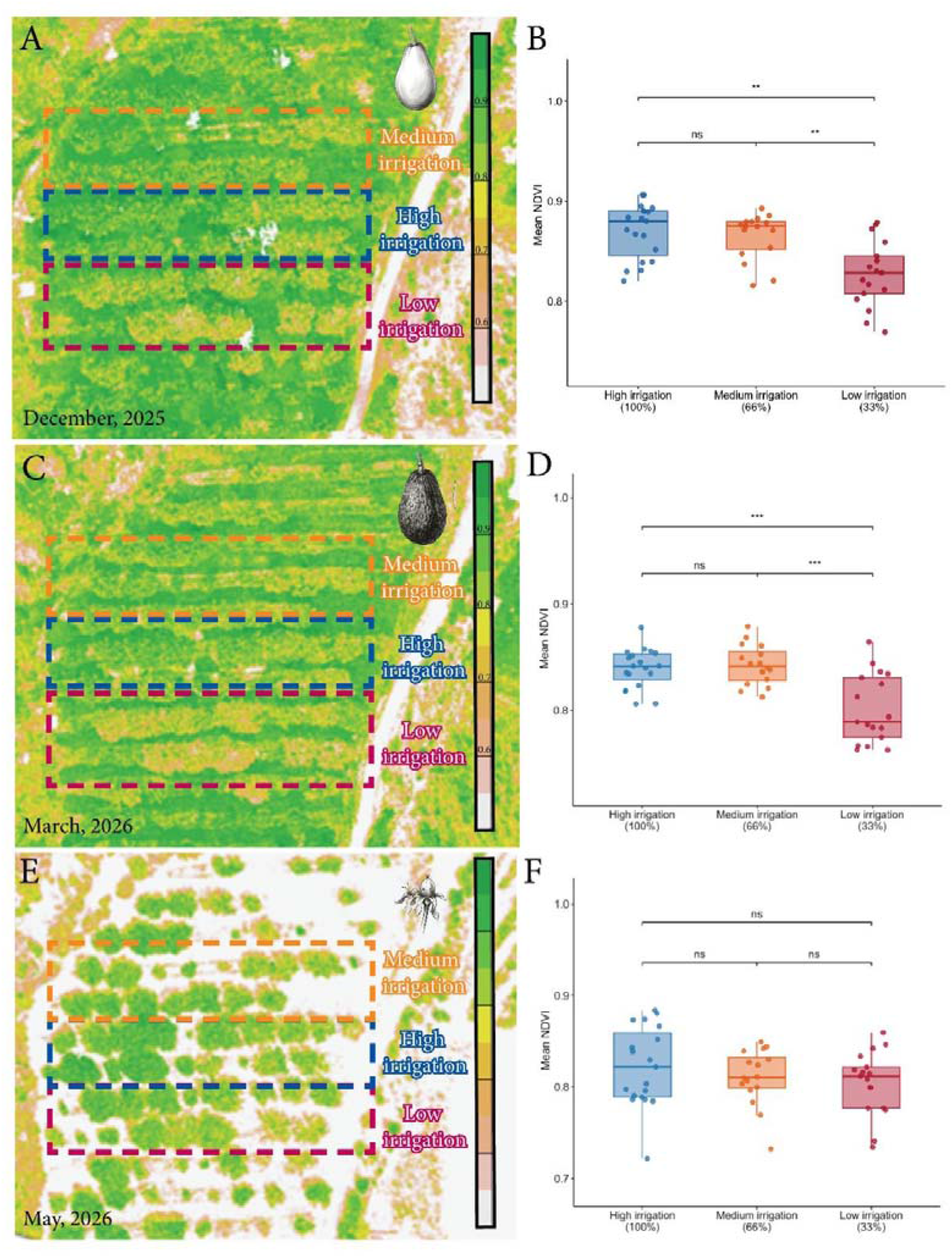
NDVI dynamics across phenology and irrigation treatments. (A) NDVI index imaged at fruit maturation in December 2025 (B) NDVI distribution by irrigation. (C) Comparison of the NDVI index at the flowering stage in March 2026. (D) NDVI distribution by irrigation. (E) NDVI index at the initial fruit set in May 2026. (F) NDVI distribution by irrigation. ANOVA test followed by a pairwise *t*-test with Holm’s correction comparing irrigation levels at *p* < 0.01(**), *p* < 0.001 (***) or “ns” (not significant).

### 3.4. Productivity and machine learning-assisted fruit weight estimation

Measurements of the fruit production revealed that the trees showed a similar number of fruits regardless of the irrigation treatment in both years 2024 (Fig. 6A) and 2025 (Fig. 6B). In contrast, while no significant differences were measured in the weight of individual fruits in 2024, those under low irrigation showed a trend towards smaller fruits (Fig. 6C). This trend became highly significant in the wet year 2025 (Fig. 6D), maintained when including the fallen fruits. Fruit drop itself was not significantly affected by the irrigation treatment, indicating that the reduction in yield under severe water deficit resulted primarily from decreased fruit size rather than increased fruit abscission.

**Figure 6.**
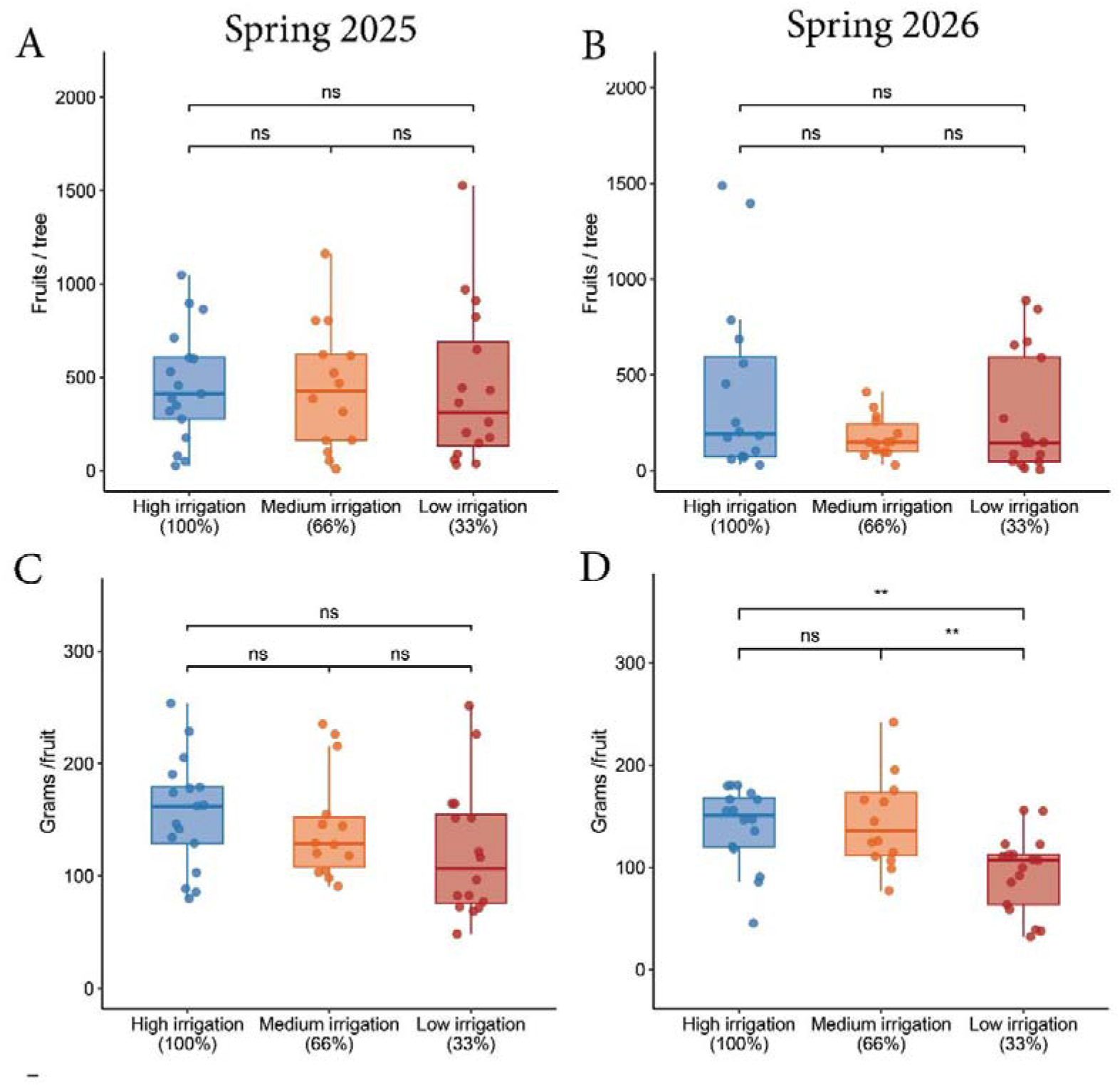
Two-year productivity of avocado during the harvest in the spring of 2025 (A, C) and the spring of 2026 (B, D). Top panels (A, B) represent the average number of fruits per tree, and bottom panels (C, D) detail the mean fruit weight. A nonparametric Kruskal-Wallis test followed by a Wilcoxon post hoc test, when significant differences were found, was used at *p* < 0.01 (**) or “ns” (not significant).

To automate weight estimation from digital imagery, a YOLOv8-seg algorithm was trained on images of fruits collected at the physiologically mature stage from all irrigation treatments. Under laboratory conditions, the model demonstrated high accuracy in fruit detection and instance segmentation, achieving a mAP@50 of 99.5% (Fig. 7A-I). Subsequently, the 2D projected area extracted from the YOLO segmentation masks was utilized to predict the physical fruit weight, yielding a strong predictive correlation (*r*^2^ = 0.83) (Fig. 7J). Moreover, this YOLO model was trained to categorize fruit according to their respective irrigation treatments. When evaluated on a separate validation dataset comprising 100 images per treatment, the algorithm successfully classified fruits from the low-, medium-, and high-irrigation treatments with accuracies of 95%, 92%, and 99%, respectively (Fig. 7K).

**Figure 7.**
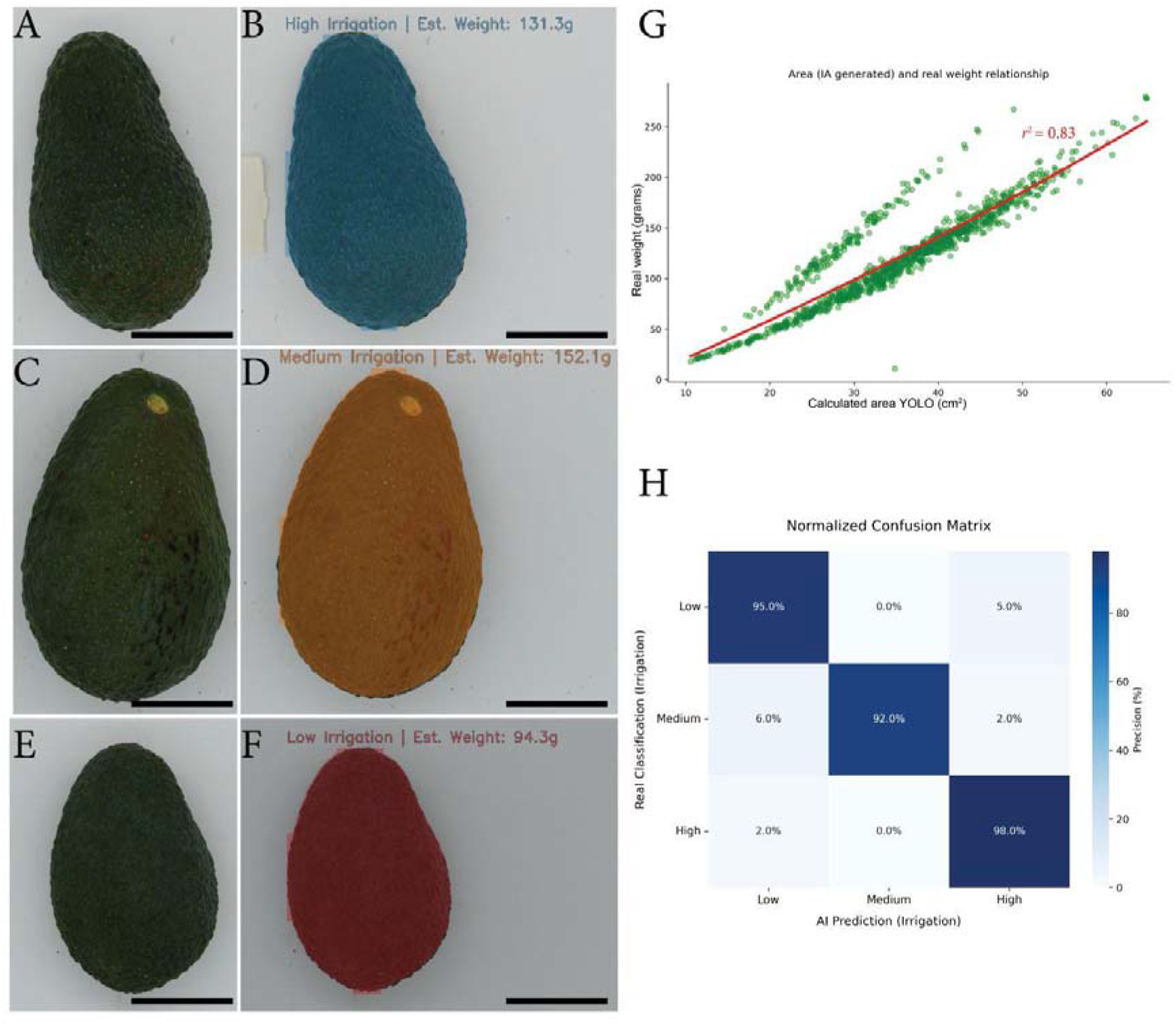
Artificial intelligence-based fruit detection and weight estimation. (A, C, E) Raw avocado images from each treatment. (B, D, F) Example YOLO segmentation and weight prediction derived from the 2D YOLO estimated area of avocado fruits per irrigation treatment. (G) Scatter plot and regression showing correlation between the estimated area of each individual fruit (green circles) calculated by YOLO (X axis, cm^2^) and the measured weight (Y axis, g). (H) Normalized confusion matrix from YOLO validation showing classification accuracy by class; values and percentages of correctly classified fruits during the YOLO model validation. A-I scale bars: 3cm.

## Discussion

This study provides a comprehensive, continuous, multi-year characterization of avocado responses to sustained deficit irrigation under Mediterranean conditions. By integrating plant-based sensors, soil moisture monitoring, remote sensing, and AI-based fruit phenotyping, this multi-scale approach revealed undocumented physiological and structural patterns that would otherwise go undetected with conventional discrete measurements. Consequently, these findings provide new insights into the mechanisms underlying avocado adaptation to prolonged water limitation while demonstrating the value of integrating complementary sensing technologies for precision irrigation management.

### Severe deficit irrigation induces a minimal physiological mode in avocado trees

This work provides highly extensive, continuous data on the physiological responses of avocado trees to sustained water scarcity under field conditions. Severe water limitation, combined with accumulated drought, markedly reduced the physiological activity of the trees. This aligns with recent studies demonstrating strict stomatal control and relatively isohydric behavior in avocado under prolonged deficit (Opazo et al., 2024; Moreno-Ortega et al., 2021). While previous works have identified xylem hydraulic embolism as a primary consequence of drought at the leaf level (Cardoso et al., 2020), our continuous monitoring reveals that both leaf thickness, a reliable integrated representation for plant water status (Afzal et al., 2017), and Ψ_trunk_ remained tightly correlated, displaying minimal diurnal fluctuations during the severe deficit irrigation regime. The pressure patch clamp has been used previously to assess water stress in several fruit trees, such as persimmon (Ballester et al., 2022; Martínez-Gimeno et al., 2017), or avocado (Rüger et al., 2010), where a strong correlation with stem and leaf water potential respectively was evidenced. Given that variations in leaf thickness are linked to stomatal conductance in avocado (Kaneko et al., 2026), this ‘physiological flattening’ (i.e., persistent stomatal closure) indicates an intense decoupling from atmospheric demand.

Furthermore, this flattening phenomenon revealed a temporal delay in the recovery process. The reduced amplitude of diurnal sensor signals persisted even as rainfall and soil water availability increased, whereas Ψ_trunk_ responded faster to rain events. This asynchronous recovery strongly points to a carry-over memory effect of accumulated drought in the soil-plant continuum (Jacques et al., 2021). The use of microtensiometers to continuously monitor the stem water potential is becoming popular in perennial fruit crops, and has been validated as a good indicator of the plant water status under certain conditions of soil water potential, such as in kiwifruit (Di Biase et al., 2025), nectarine (Conesa et al., 2023), grapevines (Lakso et al., 2022; Pagay, 2022), or citrus (Vaccaro et al., 2025), while the correlation with VPD was evidenced in pear (Blanco and Kalcsits, 2023) and apple (Blanco and Kalcsits, 2024). In avocado, the combination of continuous multi-sensor data demonstrates that 33% water irrigation accelerates leaf senescence with limited additional physiological disorders. Thus, rather than only detecting discrete stress, these results generate a novel hypothesis regarding delayed recovery and physiological legacy behaviors in perennial crops.

### Drone imaging and AI reveal novel structural and phenological parameters of drought stress in fruit crops

We initially developed a novel algorithm that measures the area of the tree canopy in real time using a recently developed methodology (Ponce et al., 2021). Tree crop phenotyping through UAV image acquisition and processing with AI has been successfully applied to other fruit crops, such as the evaluation of tree positioning and health evaluation in *Citrus* (Ampatzidis and Patel, 2019), mortality determination in oil palm (Khokthong et al., 2019), detailed crown area estimation in hazelnut (Altieri et al., 2022; Vinci et al., 2023), and crown planar area in papayas (Lai et al., 2024). While our algorithm detected tree canopy area with high accuracy, it did not detect differences across irrigation treatments in our mature avocado orchard. Among the few available studies evaluating the effects of drought on tree-crop crown size, canopy area extraction detected water stress in olive (Caruso et al., 2019, 2022). The lack of differences in our study can be attributed to the inherent architectural heterogeneity of mature avocado orchards and the limitations of 2D metrics in capturing internal canopy thinning during spring defoliation. To overcome these limitations, we evaluated the CSR, a three-dimensional descriptor that quantifies canopy surface complexity. Trees subjected to low irrigation exhibited an irregular canopy characterized by increased internal foliage gaps. While remote sensing is more commonly used to measure geometrical changes and porosity-related parameters in orchard canopies (Ponce et al., 2021), canopy gaps are specifically recognized as a meaningful structural signature of environmental disturbance (Jucker, 2021). To our knowledge, this is the first study to validate CSR as a highly sensitive remote-sensing metric of prolonged water stress in avocado, directly capturing leaf senescence and canopy thinning, which are typically triggered by chronic drought.

Furthermore, temporal NDVI analysis revealed that the spectral response of avocado trees to drought is strictly modulated the phenology. While trees under low irrigation displayed significantly lower NDVI values during fruit development than well-irrigated trees, this spectral separation disappeared during spring flowering and summer leaf flush, when NDVI values converged across all treatments. Although NDVI-based workflows are widely established for classifying plant water status in avocado orchards (Castillo-Guevara et al., 2020; Torres-Quezada et al., 2025), as well as in other fruit tree crops (Altieri et al., 2022; Vinci et al., 2023), UAV stress assessments are frequently conducted without accounting for the dynamic background of the crop’s development stages. Our continuous monitoring demonstrates that severe water deficit degrades canopy greenness at the mature developmental stages of leaves and fruits, thereby masking their spectral signatures during active vegetative flushing. This aligns with recent satellite-based observations indicating that avocado vegetation indices change significantly across vegetative growth stages (Torres-Quezada et al., 2025). Consequently, these findings emphasize that incorporating phenological context into the interpretation of remote-sensing data is essential for improving the reliability of precision irrigation systems and optimizing the management of scarce water resources.

### Integrating sensors, drones, and AI provides a holistic view of water stress boundaries in fruit crops

This work exemplifies an innovative and multi-scale approach to characterizing the spatial and temporal effects of water deprivation in mature avocado orchards. While plant water potential and stomatal conductance are widely accepted as direct indicators of water stress in avocado (Moreno-Pérez et al., 2024), traditional field assessments rely on manual, discrete measurements that are highly laborious and difficult to scale at the orchard level (Cho et al., 2024). Our 5G-enabled sensor stack successfully overcomes this constraint by capturing the rapid physiological responses of the tree continuum. The synchrony observed between continuous Ψ_trunk_ and leaf patch pressure under low irrigation conditions highlights the strict isohydric defense mechanism of this cultivar, which operates in close coordination with root-zone water dynamics across the monitored soil profile (Torres-Quezada et al., 2025). In contrast, we demonstrated that reducing irrigation to 66% has minimal effects on physiology, vegetative indices, or productivity.

While plant-based sensors only provide point-source information, they typically miss the spatial heterogeneity of mature orchards. Remote sensing is increasingly used to assess orchard productivity and predictability using satellite imagery (Benami et al., 2021). Yet, satellites are only suitable for large crop areas, limiting the technology’s potential for smaller growers and less accessible territories. Our work is pioneer in providing real-time AI-based processing and integrating it with sensors. For example, previous remote-sensing efforts have successfully linked stem water potential measurements to the field level using UAV thermal mapping (Park et al., 2017), but a multi-scale approach is still lacking. Our comprehensive spatiotemporal integration is particularly relevant for providing real-time data during critical phenological stages in crops, such as fruit development, when avocado is highly sensitive to water deficits, often leading to reduced fruit size or accelerated fruit drop (Fuentes-Peñailillo et al., 2025; Kaneko et al., 2026). By incorporating AI-based phenotyping using YOLO segmentation models, our workflow directly links atmospheric and subterranean water limitations to their ultimate productive consequences, demonstrating that long-term water deficit alters fruit weight allocation rather than total fruit count. Indeed, we provided the first approach for yield prediction in avocado under water deficit. Yield predictions, combined with phenology, have become a priority in sustainable agriculture over the last five years (Istiak et al., 2023; Bregaglio et al., 2023). We combined yield prediction with sensor networks, aerial imagery, and machine learning to provide a comprehensive framework that ultimately optimizes non-renewable resources under increasingly arid Mediterranean conditions. Recent technological support for irrigation studies suggests that the actual water requirements of Mediterranean avocado orchards can be managed significantly below conventional regional recommendations without triggering a catastrophic collapse in yield (Lilli et al., 2024). By capturing both short-term physiological dynamics and long-term structural and productive outcomes, this multi-scale study sets a robust precedent for future precision agriculture protocols in perennial fruit crops, offering actionable insights to establish precise water stress boundaries and secure sustainable crop production under climate change, a major goal in orchard and land management (Wang et al., 2025).

## 5 Conclusions

This study demonstrates that integrating continuous physiological monitoring, remote sensing, and artificial intelligence provides a comprehensive framework for precision irrigation management in mature avocado orchards. High-frequency sensing successfully captured previously undocumented hydraulic dynamics, such as the physiological flatline response under severe water deficit and drought carryover effects between seasons. Concurrently, UAV-derived CSR and phenology-modulated NDVI emerged as robust, complementary indicators for tracking long-term structural and spectral water-stress signatures. Together, these multi-scale findings prove that a unified sensor stack provides a more reliable assessment of tree water status than any individual monitoring method. Finally, beyond advancing our understanding of avocado physiological responses to water limitation, the data-driven integrated framework presented here provides a technological basis for developing plant-informed irrigation strategies that optimize water-use efficiency while ensuring crop productivity under increasingly scarce and variable water resources in Mediterranean climates.

## 6 Acknowledgments

We thank Patricia Álvarez and Vicente Capllonch for their help with fieldwork, and Javier Jiménez and Mª del Mar Moreno for their support with 5G connectivity. We also thank Pepe Ramos, Manuel Tirado, Pedro Martín, and the technical staff of IHSM-CSIC-UMA for their assistance with facility maintenance. This work was supported by the 6G-PATH project (FARM-1 use case) Grant Agreement N. 101139172, cofunded by the European Uniońs Horizon Europe research and the 6GSNS. JML was further supported by the projects PID2024-162914OB-100 from the Agencia Estatal de Investigación and a PIDI-2024-02642 from Junta de Andalucía.

## 7 Competing Interest

The authors declare that they have no known competing financial interests or personal relationships that could have appeared to influence the work reported in this paper.

## Notes

### Competing Interest Statement

The authors have declared no competing interest.

## References

Afzal, A., Duiker, S. W., & Watson, J. E. (2017). Leaf thickness to predict plant water status. Biosystems Engineering, 156, 148–156. 10.1016/j.biosystemseng.2017.01.011

Ahlmann-Eltze, C., & Patil, I. (2021). ggsignif: R Package for displaying significance brackets for’ggplot2’. PsyArxiv. 10.31234/osf.io/7awm6

Alcaraz, M. L., Hormaza, J. I., & Rodrigo, J. (2013). Pistil starch reserves at anthesis correlate with final flower fate in avocado (Persea americana). PLoS One, 8(10), e78467. 10.1371/journal.pone.0078467

Altieri, G., Maffia, A., Pastore, V., Amato, M., & Celano, G. (2022). Use of high-resolution multispectral UAVs to calculate projected ground area in Corylus avellana L. tree orchard. Sensors, 22(19), 7103. 10.3390/s22197103

Ampatzidis, Y., & Partel, V. (2019). UAV-based high throughput phenotyping in citrus utilizing multispectral imaging and artificial intelligence. Remote Sensing, 11(4), 410. 10.3390/rs11040410

Bai, T., Wang, S., Meng, W., Zhang, N., Wang, T., Chen, Y., & Mercatoris, B. (2019). Assimilation of remotely-sensed LAI into WOFOST model with the SUBPLEX algorithm for improving the field-scale jujube yield forecasts. Remote sensing, 11(16), 1945. 10.3390/rs11161945

Ballester, C., Badal, E., Bonet, L., Testi, L., & Intrigliolo, D. S. (2022). Determining transpiration coefficients of ‘Rojo Brillante’persimmon trees under Mediterranean climatic conditions. Agricultural Water Management, 271, 107804. 10.1016/j.agwat.2022.107804

Benami, E., Jin, Z., Carter, M. R., Ghosh, A., Hijmans, R. J., Hobbs, A., & Lobell, D. B. (2021). Uniting remote sensing, crop modelling and economics for agricultural risk management. Nature Reviews Earth & Environment, 2(2), 140–159. 10.1038/s43017-020-00122-y

Berry, A., Vivier, M. A., & Poblete-Echeverría, C. (2025). Evaluation of canopy fraction-based vegetation indices, derived from multispectral UAV imagery, to map water status variability in a commercial vineyard. Irrigation Science, 43(1), 135–153. 10.1007/s00271-023-00907-1

Beyá-Marshall, V., Arcos, E., Seguel, O., Galleguillos, M., & Kremer, C. (2022). Optimal irrigation management for avocado (cv.’Hass’) trees by monitoring soil water content and plant water status. Agricultural Water Management, 271, 107794. 10.1016/j.agwat.2022.107794

Blanco, V., & Kalcsits, L. (2023). Long-term validation of continuous measurements of trunk water potential and trunk diameter indicate different diurnal patterns for pear under water limitations. Agricultural Water Management, 281, 108257. 10.1016/j.agwat.2023.108257

Blanco, V., & Kalcsits, L. (2024). Relating microtensiometer-based trunk water potential with sap flow, canopy temperature, and trunk and fruit diameter variations for irrigated ‘Honeycrisp’apple. Frontiers in Plant Science, 15, 1393028. 10.3389/fpls.2024.1393028

Bregaglio, S., Ginaldi, F., Raparelli, E., Fila, G., & Bajocco, S. (2023). Improving crop yield prediction accuracy by embedding phenological heterogeneity into model parameter sets. Agricultural Systems, 209, 103666. 10.1016/j.agsy.2023.103666

Calabritto, M., Mininni, A. N., Di Biase, R., Pietrafesa, A., & Dichio, B. (2024). Spatio-temporal dynamics of root water uptake and identification of soil moisture thresholds for precision irrigation in a Mediterranean yellow-fleshed kiwifruit orchard. Frontiers in Plant Science, 15, 1472093. 10.3389/fpls.2024.1472093

Campos, J., García-Ruíz, F., & Gil, E. (2021). Assessment of vineyard canopy characteristics from vigour maps obtained using UAV and satellite imagery. Sensors, 21(7), 2363. 10.3390/s21072363

Cárceles-Rodríguez, B., Durán-Zuazo, V. H., Franco-Tarifa, D., Cuadros Tavira, S., Sacristan, P. C., & García-Tejero, I. F. (2023). Irrigation alternatives for avocado (Persea americana Mill.) in the Mediterranean subtropical region in the context of climate change: A review. Agriculture, 13(5), 1049. 10.3390/agriculture13051049

Cardoso, A. A., Batz, T. A., & McAdam, S. A. (2020). Xylem embolism resistance determines leaf mortality during drought in Persea americana. Plant physiology, 182(1), 547–554. 10.1104/pp.19.00585

Carella, A., Bulacio Fischer, P. T., Massenti, R., & Lo Bianco, R. (2024). Continuous plant-based and remote sensing for determination of fruit tree water status. Horticulturae, 10(5), 516. 10.3390/horticulturae10050516

Caruso, G., Palai, G., Tozzini, L., & Gucci, R. (2022). Using visible and thermal images by an unmanned aerial vehicle to monitor the plant water status, canopy growth and yield of olive trees (cvs. Frantoio and Leccino) under different irrigation regimes. Agronomy, 12(8), 1904. 10.3390/agronomy12081904

Caruso, G., Zarco-Tejada, P. J., González-Dugo, V., Moriondo, M., Tozzini, L., Palai, G., & Gucci, R. (2019). High-resolution imagery acquired from an unmanned platform to estimate biophysical and geometrical parameters of olive trees under different irrigation regimes. PloS one, 14(1), e0210804. 10.1371/journal.pone.0210804

Castillo-Guevara, M. A., Palomino-Quispe, F., Alvarez, A. B., & Coaquira-Castillo, R. J. (2020). Water stress analysis using aerial multispectral images of an avocado crop. In 2020 IEEE Engineering International Research Conference (EIRCON) (pp. 1–4). IEEE. 10.1109/EIRCON51178.2020.9254011

Chartzoulakis, K., Patakas, A., Kofidis, G., Bosabalidis, A., & Nastou, A. (2002). Water stress affects leaf anatomy, gas exchange, water relations, and growth of two avocado cultivars. Scientia Horticulturae, 95(1-2), 39–50. 10.1016/S0304-4238(02)00016-X

Cho, S. (2024). Integrating Cover Crops and Almond Hulls and Shells as Organic Matter Amendments for Whole Orchard Regenerative Management (Order No. 31328302). Available from ProQuest Dissertations & Theses Global. (3089712347). https://www.proquest.com/dissertations-theses/integrating-cover-crops-almond-hulls-shells-as/docview/3089712347/se-2

Conesa, M. R., Conejero, W., Vera, J., & Ruiz-Sánchez, M. C. (2023). Assessment of trunk microtensiometer as a novel biosensor to continuously monitor plant water status in nectarine trees. Frontiers in Plant Science, 14, 1123045. 10.3389/fpls.2023.1123045

Davur, Y. J., Kämper, W., Khoshelham, K., Trueman, S. J., & Bai, S. H. (2023). Estimating the ripeness of Hass avocado fruit using deep learning with hyperspectral imaging. Horticulturae, 9(5), 599. 10.3390/horticulturae9050599

Di Biase, R., Calabritto, M., Mininni, A. N., Montanaro, G., & Dichio, B. (2025). Microtensiometer-based trunk water potential as a plant water status indicator in kiwifruit under different soil water availability. Irrigation Science, 43(4), 937–954. 10.1007/s00271-025-01020-1

Durán-Zuazo, V. H., García-Tejero, I. F., Rodríguez, B. C., Tarifa, D. F., Ruiz, B. G., & Sacristán, P. C. (2021). Deficit irrigation strategies for subtropical mango farming. A review. Agronomy for sustainable development, 41(1), 13. 10.1007/s13593-021-00671-6

Durán-Zuazo, V. H., Lipan, L., Rodríguez, B. C., Sendra, E., Tarifa, D. F., Nemś, A.,…& García-Tejero, I.F. (2021). Impact of deficit irrigation on fruit yield and lipid profile of terraced avocado orchards. Agronomy for Sustainable Development, 41(5), 69. 10.1007/s13593-021-00731-x

Dwyer, B., Nelson, J., Hansen, T., et al. (2026). Roboflow (Version 1.0) [Software]. Available from https://roboflow.com. Computer vision.

Fox J and Weisberg S (2019). An R companion to applied regression, Third edition. Sage, Thousand Oaks CA. https://www.john-fox.ca/Companion.

Fuentes-Peñailillo, F., del Campo-Hitschfeld, M. L., Gutter, K., & Torres-Quezada, E. (2025). Data-driven integration of remote sensing, agro-meteorology, and wireless sensor networks for crop water demand estimation: Tools towards sustainable irrigation in high-value fruit crops. Agronomy, 15(9), 2122. 10.3390/agronomy15092122

Garrett Grolemund, Hadley Wickham (2011). Dates and times made easy with lubridate. Journal of Statistical Software, 40(3), 1–25. URL https://www.jstatsoft.org/v40/i03/.

Gutiérrez-Gordillo, S., de la Gala González-Santiago, J., Trigo-Córdoba, E., Rubio-Casal, A. E., García-Tejero, I. F., & Egea, G. (2021). Monitoring of emerging water stress situations by thermal and vegetation indices in different almond cultivars. Agronomy, 11(7), 1419. 10.3390/agronomy11071419

Harris, C. R., Millman, K. J., van der Walt, S. J., Gommers, R., Virtanen, P., Cournapeau, D.,…& Oliphant, T. E. (2020). Array programming with NumPy. Nature, 585, 357–362. 10.1038/s41586-020-2649-2

Hijmans, R. (2025). _terra: Spatial Data Analysis_. R package version 1.8-50, https://CRAN.R-project.org/package=terra.

Houetohossou, S. C. A., Houndji, V. R., Hounmenou, C. G., Sikirou, R., & Kakaï, R. L. G. (2023). Deep learning methods for biotic and abiotic stresses detection and classification in fruits and vegetables: State of the art and perspectives. Artificial Intelligence in Agriculture, 9, 46–60. 10.1016/j.aiia.2023.08.001

Hunter, J. D. (2007). Matplotlib: A 2D graphics environment. Computing in Science & Engineering, 9(3), 90–95. 10.1109/MCSE.2007.55

Istiak, M. A., Syeed, M. M., Hossain, M. S., Uddin, M. F., Hasan, M., Khan, R. H., & Azad, N. S. (2023). Adoption of Unmanned Aerial Vehicle (UAV) imagery in agricultural management: A systematic literature review. Ecological Informatics, 78, 102305. 10.1016/j.ecoinf.2023.102305

Jacques, C., Salon, C., Barnard, R. L., Vernoud, V., & Prudent, M. (2021). Drought stress memory at the plant cycle level: A review. Plants, 10(9), 1873. 10.3390/plants10091873

Jucker, T. (2022). Deciphering the fingerprint of disturbance on the three dimensional structure of the world’s forests. New Phytologist, 233(2), 612–617. 10.1111/nph.17729

Junquera, V., Hormaza, J. I., Rubenstein, D. I., Levin, S. A., Vadillo Pérez, I., & Gavilán, P. J. (2025). Severe water crisis in southern Spain under expanding irrigated agriculture: A multidimensional drought analysis. Proceedings of the National Academy of Sciences, 122(39), e2508055122. 10.1073/pnas.2508055122

Kaneko, T., Gould, N., Campbell, D., & Clearwater, M. J. (2026). Irrigation, water deficit and crop load effects on ‘Hass’ avocado fruit size under New Zealand growing Conditions. Horticulturae, 12(2), 230. 10.3390/horticulturae12020230

Khokthong, W., Zemp, D. C., Irawan, B., Sundawati, L., Kreft, H., & Hölscher, D. (2019). Drone-based assessment of canopy cover for analyzing tree mortality in an oil palm agroforest. Frontiers in Forests and Global Change, 2, 12. 10.3389/ffgc.2019.00012

Lai, S., Ming, H., Huang, Q., Qin, Z., Duan, L., Cheng, F., & Han, G. (2024). Remote sensing extraction of crown planar area and plant number of papayas using UAV images with very high spatial resolution. Agronomy, 14(3), 636. 10.3390/agronomy14030636

Lakso, A. N., Santiago, M., & Stroock, A. D. (2022). Monitoring stem water potential with an embedded microtensiometer to inform irrigation scheduling in fruit crops. Horticulturae, 8(12), 1207. 10.3390/horticulturae8121207

Lilli, M. A., Efstathiou, D., Koukianaki, E. A., Paranychianakis, N., & Nikolaidis, N. P. (2024). Optimizing the water-ecosystem-food nexus of avocado plantations. Frontiers in Water, 6, 1412146. 10.3389/frwa.2024.1412146

Longchamps, L., Tisseyre, B., Taylor, J., Sagoo, L., Momin, A., Fountas, S., & Khosla, R. (2022). Yield sensing technologies for perennial and annual horticultural crops: a review. Precision Agriculture, 23(6), 2407–2448. 10.1007/s11119-022-09906-2

Magh, R. K., Paligi, S. S., Papastefanou, P., Klosterhalfen, A., Ammer, C., Beyer, M.,…& Hildebrandt, A. (2026). Continuous stem water potential measurements with microtensiometry reveal species identity and soil matric potential control of stem water potential in temperate forests. Ecohydrology, 19(3), e70197. 10.1002/eco.70197

Marino, G., Scalisi, A., Guzmán-Delgado, P., Caruso, T., Marra, F. P., & Lo Bianco, R. (2021). Detecting mild water stress in olive with multiple plant-based continuous sensors. Plants, 10(1), 131. 10.3390/plants10010131

Martínez-Gimeno, M. A., Castiella, M., Rüger, S., Intrigliolo, D. S., & Ballester, C. (2017). Evaluating the usefulness of continuous leaf turgor pressure measurements for the assessment of Persimmon tree water status. Irrigation Science, 35(2), 159–167. 10.1007/s00271-016-0527-3

Massaad, M., Scuderi, D., Andolina, F., Bellitti, S., Buscaglia, A., Gugliuzza, G., & Farina, V. (2026). Tropical fruits in the Mediterranean Basin: current research status, priorities, and knowledge gaps. *A systematic review*. Frontiers in Plant Science, 17, 1817537. 10.3389/fpls.2026.1817537

McKinney, W. (2010). Data structures for statistical computing in Python. Proceedings of the 9th Python in Science Conference, 56–61. 10.25080/Majora-92bf1922-00a

Miranda, J. C., Arnó, J., Gené-Mola, J., Lordan, J., Asín, L., & Gregorio, E. (2023). Assessing automatic data processing algorithms for RGB-D cameras to predict fruit size and weight in apples. Computers and Electronics in Agriculture, 214, 108302. 10.1016/j.compag.2023.108302

Moreno-Ortega, G., Pliego, C., Sarmiento, D., Barceló, A., & Martínez-Ferri, E. (2019). Yield and fruit quality of avocado trees under different regimes of water supply in the subtropical coast of Spain. Agricultural Water Management, 221, 192–201.10.1016/j.agwat.2019.05.001

Moreno-Ortega, G., Zumaquero, A., Matas, A., Olivier, N. A., van den Berg, N., Palomo-Ríos, E., Martínez-Ferri, E., & Pliego, C. (2021). Physiological and molecular responses of ‘Dusa’ avocado rootstock to water stress: Insights for drought adaptation. Plants, 10(10), 2077. 10.3390/plants10102077

Moreno-Pérez, A., Barceló, A., Pliego, C., & Martínez-Ferri, E. (2024). Water relations and physiological response to water deficit of ‘Hass’ avocado grafted on two rootstocks tolerant to *R. necatrix*. Agronomy, 14(9), 1959. 10.3390/agronomy14091959

Nemera, D. B., Bar-Tal, A., Levy, G. J., Tarchitzky, J., Rog, I., Klein, T., & Cohen, S. (2021). Mitigating negative effects of long-term treated wastewater irrigation: Leaf gas exchange and water use efficiency response of avocado trees (*Persea americana* Mill.). Agricultural Water Management, 256, 107126. 10.1016/j.agwat.2021.107126

Opazo, I., Pimentel, P., Salvatierra, A., Ortiz, M., Toro, G., & Garrido-Salinas, M. (2024). Water stress tolerance is coordinated with water use capacity and growth under water deficit across six fruit tree species. Irrigation Science, 42(3), 493–507. 10.1007/s00271-024-00915-9

Ortuño, M. F., Conejero, W., Moreno, F., Moriana, A., Intrigliolo, D. S., Biel, C.,…& Torrecillas, A. (2010). Could trunk diameter sensors be used in woody crops for irrigation scheduling? A review of current knowledge and future perspectives. Agricultural Water Management, 97(1), 1–11. 10.1016/j.agwat.2009.09.008

Pagay, V. (2021). Dynamic aspects of plant water potential revealed by a microtensiometer. BioRxiv, 2021-06. 10.1101/2021.06.23.449675

Pagay, V. (2022). Evaluating a novel microtensiometer for continuous trunk water potential measurements in field-grown irrigated grapevines. Irrigation Science, 40(1), 45–54. 10.1007/s00271-021-00758-8

Park, S., Ryu, D., Fuentes, S., Chung, H., Hernández-Montes, E., & O’Connell, M. (2017). Adaptive estimation of crop water stress in nectarine and peach orchards using high-resolution imagery from an unmanned aerial vehicle (UAV). Remote Sensing, 9(8), 828. 10.3390/rs9080828

Ponce, J. M., Aquino, A., Tejada, D., Al-Hadithi, B. M., & Andújar, J. M. (2021). A methodology for the automated delineation of crop tree crowns from UAV-based aerial imagery by means of morphological image analysis. Agronomy, 12(1), 43. 10.3390/agronomy12010043

R Core Team (2023) R: A Language and environment for statistical computing. R Foundation for statistical computing, Vienna. https://www.R-project.org/

Robson, A., Rahman, M. M., & Muir, J. (2017). Using worldview satellite imagery to map yield in avocado (*Persea americana*): A case study in Bundaberg, Australia. Remote Sensing, 9(12), 1223. 10.3390/rs9121223

Rodríguez-Domínguez, C. M., Hernandez-Santana, V., Buckley, T. N., Fernández, J. E., & Díaz-Espejo, A. (2019). Sensitivity of olive leaf turgor to air vapour pressure deficit correlates with diurnal maximum stomatal conductance. Agricultural and Forest Meteorology, 272, 156–165. 10.1016/j.agrformet.2019.04.006

Roussel, J.R., Auty, D., Coops, N. C., Tompalski, P., Goodbody, T. R. H., Sánchez Meador, A., Bourdon, J.F., De Boissieu, F., Achim, A. (2020). lidR: An R package for analysis of Airborne Laser Scanning (ALS) data. Remote Sensing of Environment, 251 (August), 112061. 10.1016/j.rse.2020.112061.

Rüger S., Ehrenberger W., Arend M., Geßner P., Zimmermann G., Zimmermann D., Bentrup F.W., Nadler A., Raveh E., Sukhorukov V.L., Zimmermann U. (2010). Comparative monitoring of temporal and spatial changes in tree water status using the non-invasive leaf patch clamp pressure probe and the pressure bomb. Agriculture Water Management, 98, 283–290. 10.1016/j.agwat.2010.08.022.

Sharma, K., & Shivandu, S. K. (2024). Integrating artificial intelligence and Internet of Things (IoT) for enhanced crop monitoring and management in precision agriculture. Sensors International, 5, 100292. 10.1016/j.sintl.2024.100292

Torres-Quezada, E., Fuentes-Peñailillo, F., Gutter, K., Rondón, F., Marmolejos, J. M., Maurer, W., & Bisono, A. (2025). Remote sensing and soil moisture sensors for irrigation management in avocado orchards: A practical approach for water stress assessment in remote agricultural areas. Remote Sensing, 17(4), 708. 10.3390/rs17040708

Vaccaro, G., Fusco, M., Alagna, V., Franco, L., Motisi, A., & Iovino, M. (2025). Assessing microtensiometers for monitoring stem water potential in mandarin (*Citrus reticulata* Blanco) orchard under different irrigation regimes. Agricultural Water Management, 320, 109873. 10.1016/j.agwat.2025.109873

Velazquez-Chavez, L. J., Daccache, A., Mohamed, A. Z., & Centritto, M. (2024). Plant-based and remote sensing for water status monitoring of orchard crops: Systematic review and meta-analysis. Agricultural Water Management, 303, 109051. 10.1016/j.agwat.2024.109051

Vinci, A., Brigante, R., Traini, C., & Farinelli, D. (2023). Geometrical characterization of hazelnut trees in an intensive orchard by an unmanned aerial vehicle (UAV) for precision agriculture applications. Remote Sensing, 15(2), 541. 10.3390/rs15020541

Wang, X., Li, Z., Li, H., Shen, T., Luo, Y., Zhang, F.,…& Zhang, X. (2025). Maize straw application shows regional-scale improvements to soil fertility and crop yields in Chinese croplands: A meta-analysis. Field Crops Research, 333, 109908. 10.1016/j.fcr.2025.109908

Wheeler, W.D, Black, B. & Bugbee, B. (2023) Assessing water stress in a high-density apple orchard using trunk circumference variation, sap flow index and stem water potential. Frontiers in Plant Science. 14:1214429. 10.3389/fpls.2023.1214429

Wickham, H., François, R., Henry, L., Müller, K., Vaughan, D. (2023). _dplyr: A Grammar of Data Manipulation. R package version 1.1.4, https://CRAN.R-project.org/package=dplyr.

Wickham, H., & Bryan, J. (2023). R packages. O’Reilly Media.

Yin, H., Cao, Y., Marelli, B., Zeng, X., Mason, A. J., & Cao, C. (2021). Soil sensors and plant wearables for smart and precision agriculture. Advanced Materials, 33(20), 2007764. 10.1002/adma.202007764

